# Prior-guided Individualized Thalamic Parcellation based on Local Diffusion Characteristics using Deep Learning

**DOI:** 10.1101/2022.07.19.500596

**Authors:** Chaohong Gao, Xia Wu, Yaping Wang, Gang Li, Congying Chu, Kristoffer Hougaard Madsen, Lingzhong Fan

## Abstract

As a gateway for projections entering and exiting the cerebral cortex, the human thalamus processes information from sensory to cognition relevant to various neuropsychiatric disorders. It is composed of dozens of nuclei, which have been difficult to identify with clinical MR sequences. However, delineating thalamic nuclei accurately at an individual level is essential for precise neuromodulation treatment. Here, we not only identified the fine-grained thalamic nuclei using local diffusion properties in vivo but also employed a deep learning strategy to achieve highly reproducible individual-level parcellation. Using High-quality diffusion MRI (dMRI), we first constructed a fine-grained group thalamus atlas based on thalamic local diffusion features. Then, the high-probability core area of the group thalamus atlas was wrapped into the native space as prior guidance for individualized thalamus construction. Finally, we trained the semi-supervised multiple classification models to accurately construct the individualized thalamus atlas with single-subject local diffusion characteristics. Compared to group atlas registration and single-subject clustering strategies, our individualized thalamus atlas combines population commonality and individual specificity and is superior in depicting the individual thalamic nuclei boundaries. Besides, our atlas provides a more conspicuous capacity to capture the individual specificity of thalamic nuclei. Through the evaluation by 3.0T\7.0T and test-retest dMRI datasets, the proposed high-probability group prior guided individualized thalamus construction pipeline is robust and repeatable in different magnetic field strengths and scanning batches. In addition, the individual parcellation of the thalamic nuclei has a good correspondence with the histological atlas and captured both higher group consistency and inter-subjects variations, which could be a valuable solution for precision clinical treatment.

## Introduction

The dorsal thalamus, usually called the thalamus, is the largest part of the diencephalon, which can be divided generally into anterior, lateral, and medial nuclei by internal medullary lamina ^1^. These nuclei are not only differentiated in structure but also responsible for different functions ^2–4^, and have also been shown to be selectively vulnerable to specific pathological conditions ^5^. Therefore, in recent years, specific thalamic nuclei have become the neuromodulatory target of nervous system diseases. For example, deep brain stimulation of the ventralis intermedius nucleus is effective in inhibiting essential tremors ^6^, and the anterior thalamic nucleus is the target of intractable epilepsy ^7^, the ventral intermediate thalamic nucleus to treat Parkinson’s disease and essential tremor ^8^; the ventral posterolateral nucleus to relieve neuropathic pain ^9^, the centromedian-parafascicular-complex for consciousness recovery after severe traumatic brain injury ^10^ as well as Tourette Syndrome ^11^. However, the therapeutic effect differs in patients, which indicates that the thalamic structure of each individual still has particular specificity, especially in patients with specific diseases.

The human thalamic nuclei play varying roles in different brain disorders, but the boundaries of each thalamus nucleus can only be directly traced by the histological atlases ^12–15^. Although histology is usually considered the gold standard, the experience of different anatomists, different histological staining methods, and the limited availability of the used specimens have led to the lack of a consistent and unified understanding of the anatomical delineation of the thalamic nuclei. Recently, the MRI-based thalamus atlas can noninvasively extract the structural or functional characteristics of the thalamus to make the thalamus atlas automatically and reliably. The imaging-based parcellation of the thalamus can be categorized into three types regarding different MRI modalities: structural MRI ^16–18^, diffusion MRI (dMRI) ^19–25^, and functional MRI ^26–29^ Parcellations based on structural MRI relied on manual labeling, and parcellations based on diffusion MRI or functional MRI were data-driven methods using the local information or global connectivity information ^26,28,30–35^. While the functional connectivity (FC) and structural connectivity (SC) can provide convincing thalamus parcellations, they have ignored the local diffusion features of the thalamus, which make more sense to depict the thalamic microstructure ^25^. To reconstruct the local diffusion pattern in the human brain, both diffusion tensor ^36–38^ and orientation distribution function (ODF) ^39–44^ based model have been proposed. Compared to DTI sampling and grid sampling in q-space, the high angular resolution diffusion imaging (HARDI) can more effectively and accurately collect the diffusion signal by shell sampling ^45^. Thanks to the HARDI techniques, the high-order ODF model can be estimated to depict detailed diffusion patterns of brain tissue with multi-shell multi-tissue constrained spherical deconvolution (MSMT-CSD) ^44^. The feasibility of constructing the thalamus atlas in a coarser granularity using ODF has been demonstrated in recent years ^25,46,47^.

Furthermore, most atlases still rely on the registration method to map themselves to individuals, assuming that nuclei are invariant in size, shape, or position between individuals. However, these differences between individuals indeed exist. Obtaining a thalamus atlas at the individual level is a critical step toward understanding the stability of the association between structure and function in the human brain across individuals ^48^. Meanwhile, the capacity to identify the unique architecture of an individual’s thalamus is essential for personalized medicine. The individual-specific brain atlas could provide higher parcellation accuracy ^49,50^ and better predict performance in cognition ^51^, behavior ^52^, and brain aging ^53^. Besides, an individualized brain atlas has been frequently applied in disease diagnosis ^54^, neurosurgery localization ^55–57^, and prognosis prediction ^58^. However, few individualized atlas construction studies focus on mapping the thalamus, especially based on the local diffusion features. Thus, developing a reliable individualization method for thalamus atlas construction is urgent and essential.

Many studies ^50,54,59–62^ have emphasized the significance of constructing the individualized brain atlas. The existing brain atlas individualization techniques are single-subject based or multi-subject based. Single-subject-based individualization relies on high-quality diffusion MRI data to reconstruct fiber projections between ROI and cerebral cortex ^24,57,63,64^, or robust parcellation algorithms to assign voxels into corresponding clusters ^52,60,65^. And they lack canonical cluster labeling criteria and population commonality. Multi-subject-based methods first build a group-level brain atlas to substitute ^62,66^ or guide ^24,50,53,67^ the further construction of an individualized atlas. The group substitution strategy completely ignores individual specificity. In contrast to these two individualization methods, group prior guided individualized atlas construction aims to combine the population commonality and individual specificity. But the combination strategy remains an unsolved issue because of the unavoidable error in the registration between group prior and personal features. To solve this, we ingeniously registered the high-probability core area of the group atlas to native space as the prior guidance. The deep learning model has flourished in neuroscience ^68^ for its robust data fitting and classification performance. And our high-probability group prior guided brain atlas individualization strategy can be considered a kind of semi-supervised classification.

In the current study, we built a human thalamus atlas using a large population data set and the well-scanned diffusion MRI and proposed a deep learning approach to personalize it to individuals based on a high-probability group prior guided individualization strategy. Here, based on local diffusion features of thalamic nuclei, we first constructed a canonical group thalamus atlases as the prior guidance, then built the individualized thalamus atlas by the high-probability group prior guidance and deep-learning-based semi-supervised classification. We have constructed local diffusion features based group thalamus atlas with tinier granularity. Besides, tested by four high-quality data sets, our thalamus atlas individualization pipeline has been proven more powerful in capturing individual specificity than single-subject clustering and group atlas registration methods. And at the more refined parcellation granularity level, our individualization pipeline can provide even better performance than coarse parcellation numbers. Through 3.0T-7.0T and test-retest examination, this pipeline is robust to different magnetic field strengths and scanning batches. Examined by Allen brain data set, our prior-guided individualized thalamus atlas is most histologically consistent compared to single-subject clustering and group atlas registration.

## Materials and Methods

### Data Acquisition

The data sets used in this paper come from Human Connectome Project ^69^ (HCP) (https://humanconnectome.org). T1w MRI, 3T dMRI and 7T dMRI data in HCP S1200 were used. Table 1 is the profile of the dMRI data used in this paper. According to the usage, we extracted five subsets from HCP S1200. We named them as follows, HCP Unrelated 100 (N=100, 3T dMRI) data set was used to construct the group thalamus atlas and HCP 3T (N=100, 3T dMRI), HCP 7T (N=100, 7T dMRI), HCP Test (N=30, 3T dMRI), HCP Retest (N=30, 3T dMRI) data sets were to construct the individualized thalamus atlas. The HCP 3T and HCP 7T data sets have the same subject list. The HCP Test and HCP Retest have the same subject list. The subject lists of these four data sets have no overlap with HCP Unrelated 100. The subject information can be seen in the supplementary sheet 1.

**Table 1.** Profile of the dMRI data used in this paper.

|  | HCP 3T dMRI | HCP 7T dMRI |
| --- | --- | --- |
| Scan Sequence | SE-EPI | SE-EPI |
| TR (*ms*) | 5520 | 7000 |
| TE (*ms*) | 89.5 | 71.2 |
| Flip Angle (°) | 78 | 90 |
| Resolution (*mm*<sup>3</sup>) | 1.25 × 1.25 × 1.25 | 1.05 × 1.05 × 1.05 |
| *b* Value (*s/mm*<sup>2</sup>) | 1000,2000,3000 | 1000,2000 |
| Volume | 276 (6 + 270) | 136 (6 + 130) |

### Data Preprocessing

#### ROI definition

The HCP S1200 data set has been preprocessed by the HCP minimal preprocessing pipeline ^70^. Morel’s histological thalamus atlas ^13^ was selected as the initial ROI and registered to native diffusion space by FSL ^71^. As seen in S.1(a), we first linearly registered the diffusion space (B0) to structural space (T1w) and nonlinearly registered structural space to MNI space ^72^. The registration matrix was then generated by converting and inversing these two deformation fields. As seen in S.1(b), to remove the white matter (WM) and cerebrospinal fluid (CSF), the fractional anisotropy (FA) map was calculated by FSL, and the CSF probability map was calculated by SPM ^73^. Finally, in native diffusion space, voxels in thalamus ROI with FA value higher than 0.6 or CSF probability higher than 0.05 were removed.

#### ODF estimation

As shown in S.2, the ODF estimation was performed in MRtrix3 ^74^. The HCP 3T dMRI and HCP 7T dMRI data have more than 2 different non-zero *b* factors diffusion-weighted scanning. Thanks to this, the *dhollander*^75,76^ algorithm was used to compute the response function, and the MSMT-CSD ^44^ method was applied to estimate the 8th-order ODFs of thalamic voxels. Thus, the local diffusion features of each voxel were depicted by 45-dimensional spherical harmonic (SH) function coefficients.

### Group Thalamus Atlas Construction

#### Individual parcellation construction

Spectral clustering ^77^ was used to cluster thalamic voxels that close in diffusion patterns and spatial locations. Therefore, the similarity between voxels was calculated based on the 45-dimensional SH coefficients and 3-dimensional spatial location. The similarity coefficient between a voxel pair was calculated by equation (1).

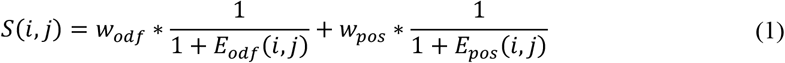

In this equation, *S*(*i,j*) is the similarity coefficient between voxel *i* and *j*; *E_odf_*(*i,j*) is the Euclidean distance of the SH coefficients, and *E_pos_*(*i,j*) is the Euclidean distance of the spatial coordinates; *w_odf_* and *w_pos_* are the weighted coefficients of the SH distance and spatial distance respectively. In this paper, these two weighted coefficients were equally set to 0.5. To avoid the bias in clustering ^25^, we scaled the SH coefficients by multiplying a scaling factor of 89 when processing HCP 3T dMRI data and 98 when 7T. To find the optimal cluster number, we set the maximum cluster number as 28, which is four times larger than the previous study ^25^.

#### Group probability atlas construction

The cluster labels among individual parcellations did not follow the same labeling criteria, making it challenging to generate the group atlas. To solve this, the individual parcellation was registered from the native diffusion space to the MNI space. An N-dimensional (N=100) label vector was generated to represent the voxel-wise labeling pattern for each voxel in ROI. The group labeling criteria could be developed in the MNI space by computing the labeling pattern similarity of all voxels and performing spectral clustering again. Then, based on the principle of maximum spatial overlap ^78^, the cluster labels could be uniformly and symmetrically rewritten. The group probability map (PM) of the thalamus was further generated by calculating the probability of voxels belonging to corresponding clusters among the individual parcellations ^23^. Finally, we set a probability threshold (0.25) on the probability map, and the survived voxels were assigned to the maximum probability cluster to construct the maximum probability map (MPM). Fig. 1 shows the whole pipeline for group thalamus atlas construction.

**Fig. 1.**
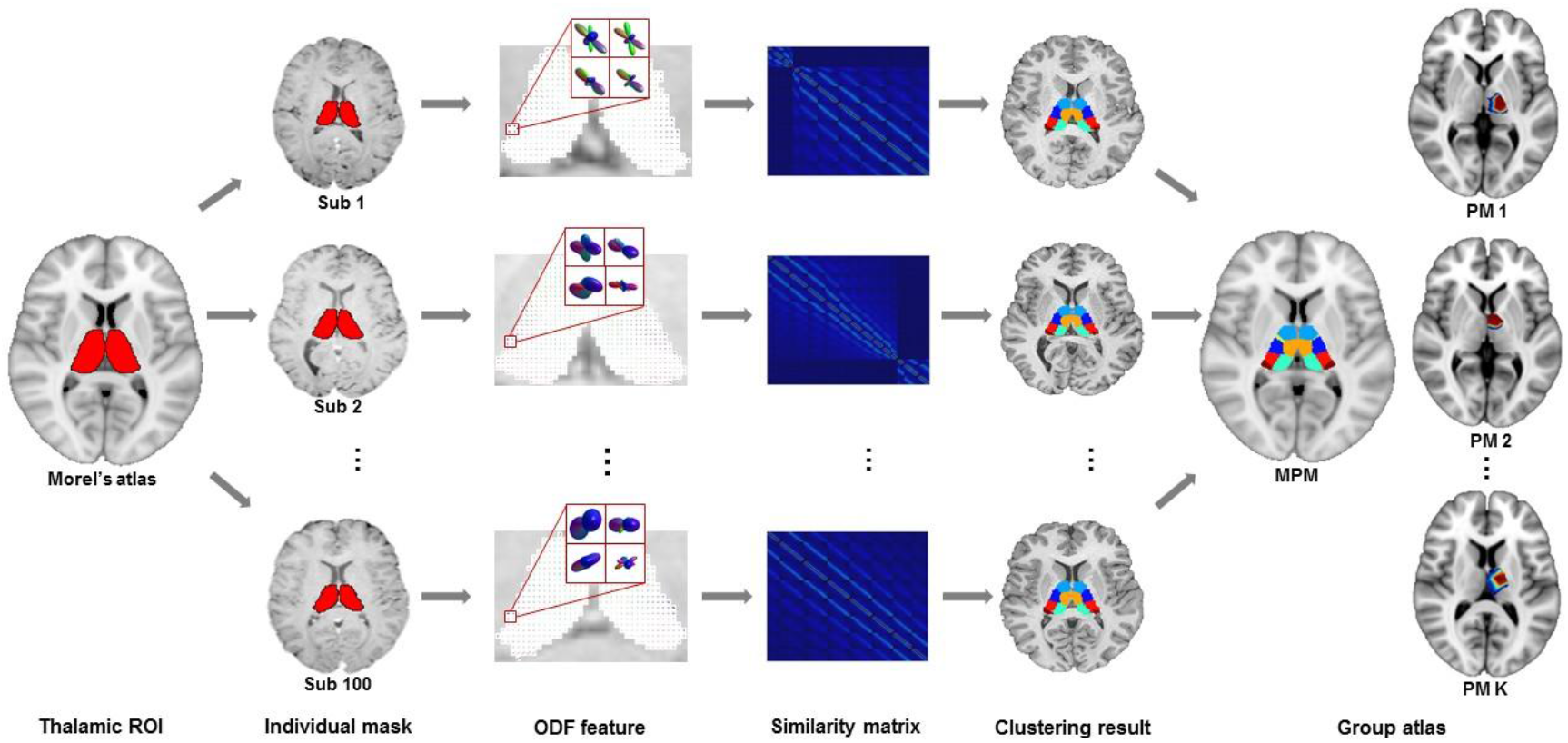
Pipeline of group thalamus atlas construction. There are several steps to construct the group thalamus atlas, including ROI definition, ROI registration and postprocessing, ODF estimation, similarity matrix calculation, spectral clustering, cluster relabeling, and probability map construction.

#### Optimal cluster numbers validation

Two validation indexes were computed to confirm the optimal cluster numbers of MPM. Dice coefficient ^79^ (*Dice*) depicts the spatial consistency between the individual atlas and group atlas. We applied a leave-one-out strategy to divide the HCP Unrelated 100 data set into 1-subject and 99-subject data sets. The mean Dice coefficient of 100 times leave-one-out validation was computed for each cluster number. Topological distance coefficient ^80^ (*TpD*) was to depict the topological consistency between two hemispheres of the group thalamus atlas. Local higher *Dice* and lower *TpD* indicate the hidden optimal cluster numbers of thalamus atlas ^23^.

### Individualized Thalamus Atlas Construction

#### Group prior guided individualized atlas construction

It has been proved that a high probability group atlas is the commonality among population ^50,53^. Based on this, we directly registered the high probability (P>0.70) group thalamus atlas to native diffusion space and took it as the high-probability group prior. Then, the construction of an individualized atlas could be regarded as semi-supervised multiple classification task. The training and testing sets were the group prior atlases and thalamic voxels outside the prior atlas (undefined atlas). The features of each voxel were 45-dimensional SH coefficients and 3-dimensional coordinates. The labels were the K-dimensional probability vector corresponding to K clusters. And the cluster label with maximum probability was assigned to the testing voxel. Each subject would receive a unique deep learning model for individualized atlas construction. The final individualized thalamus atlas was constructed by combining prior atlas and tested atlas. The whole pipeline of individualized thalamus atlas construction can be seen in Fig. 2.

**Fig. 2.**
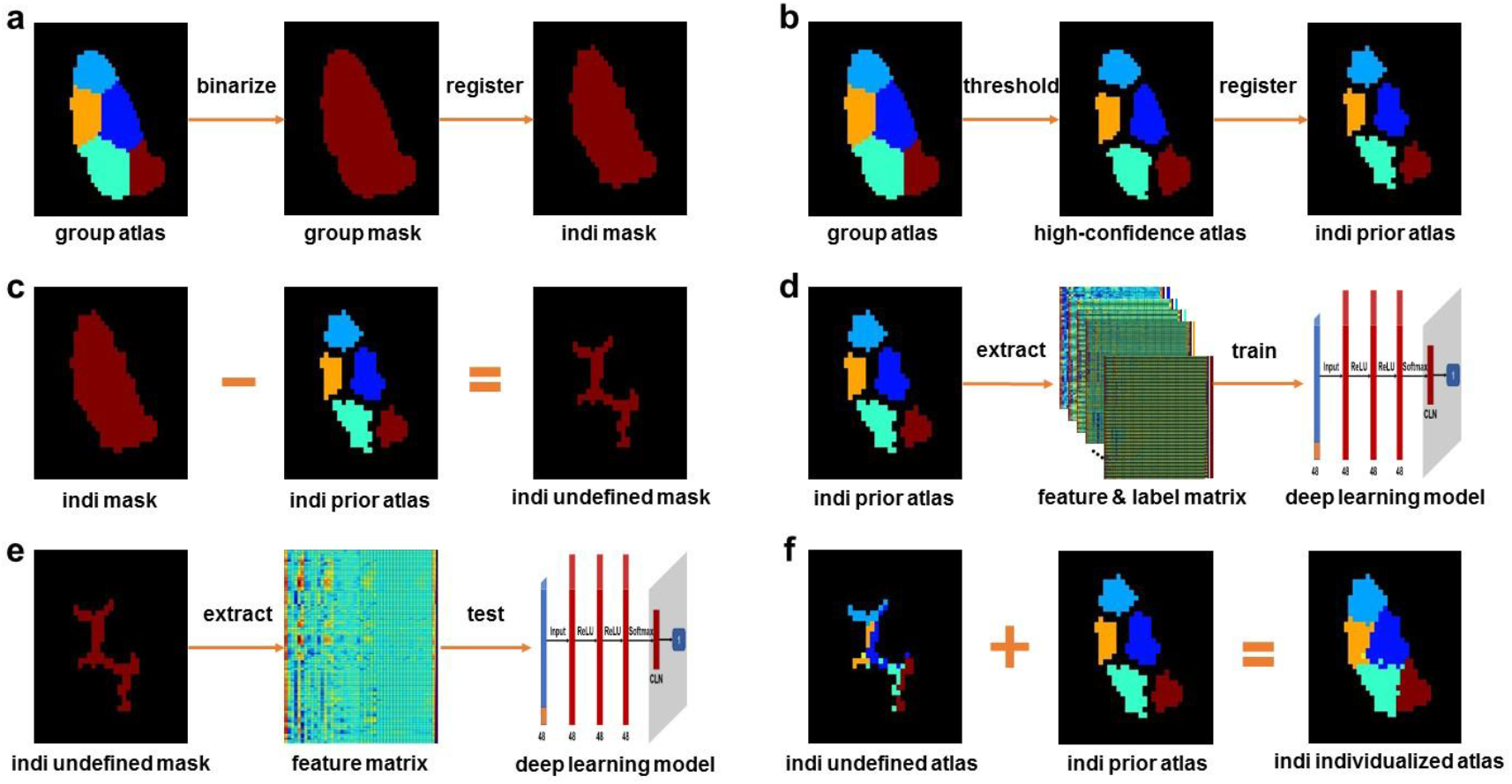
Pipeline of individualized thalamus atlas construction. (a) Binarize the group thalamus atlas with a moderate probability threshold (0.25) to generate the group mask and register it to the native space. (b) Filter the group atlas with a high probability threshold (P>0.70) to generate a high-probability group prior atlas in native space by registration. (c) Obtain the individual undefined mask by subtracting the voxels in the prior atlas from the individual mask. (d) Extract features and labels from prior atlas and train the multiple classifiers based on the deep learning model. (e) Extract features from undefined voxels and predict the probability belonging to corresponding clusters. Maximum probability cluster labels are assigned to undefined voxels and generate the undefined atlas. (f) Construct the individualized thalamus atlas by merging the prior and undefined atlas.

#### Validation of individualized thalamus atlas

To ensure the population commonality was accurately registered to each subject, the probability thresholds from 70% to 100% were adopted to generate the high-confidence group prior atlas. In addition, to check the effectiveness and generalization ability of the individualization model of each subject, a 10-fold cross validation was conducted. Cohesion within and separation between clusters is the criteria for evaluating the clustering goodness ^81^. Here, we quantified the clustering goodness by silhouette coefficient ^23,81^ (*Sil*) in equation (2).

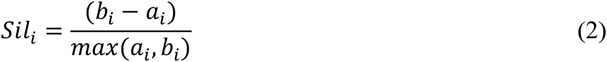

In this equation, *a_i_* represents the mean distance between voxel *i* and voxels in the same cluster; *b_i_* represents the mean distance between voxels *i* and voxels in other clusters. The distance measurement was the Euclidean distance of 45-dimensional SH coefficients, which directly depicted the local diffusion features of thalamic nuclei. A higher silhouette coefficient indicates better clustering goodness. To evaluate our group prior guided (PG) individualized thalamus atlas, we checked the clustering goodness of the individualized thalamus atlas generated by single-subject clustering (CL) and group atlas registration (GR). The single-subject clustering was to construct the individualized thalamus atlas by performing spectral clustering on the similarity matrix (the same generation pipeline in group atlas construction). The group atlas registration method was to directly register the MPM to the native space and regard it as the individualized thalamus atlas of the subject.

Besides, we quantitatively compared the improvement of clustering goodness between single-subject-based individualization and multi-subject-based. The improvement of clustering goodness (*Imp*) was computed by equation (3).

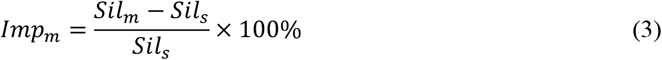

In this equation, *Sil_m_* is the silhouette coefficient of PG or PR individualized thalamus atlas, and *Sil_s_* is the CL one. The clustering goodness, as well as the improvement of it, was calculated on each hemisphere independently. We calculated these two validation indexes on HCP 3T, HCP 7T, HCP Test, and HCP Retest data sets to check the robustness of the individualization pipeline.

In addition, we checked the histological consistency by Allen brain data set ^82^. Using the MRI data provided by (atlas.brain-map.org), we constructed the individualized thalamus atlases based on prior-guided, group registration and single-subject clustering strategies. Then, the histological consistency could be examined by visually comparing the individualized thalamus atlases with manually labeled thalamus atlas by Nissl.

## Result

### Group thalamus atlas

We developed an automatic group thalamus atlas construction pipeline and constructed the probability map and maximum probability map of the thalamus based on the HCP Unrelated 100 data set. To qualify spatial consistency between subjects and homology between hemispheres, the Dice coefficient, topological distance coefficient, and topological distance coefficient were computed for cluster numbers varying from 2 to 28. According to authoritative judgment criteria ^23^, the intrinsic optimal cluster numbers were embedded in the local extreme of these two validation indexes. In Fig. 3, when the cluster number is 7 and 12, the *Dice* coefficient shows acceptable local maxima in both left and right hemispheres, and the *TpD* coefficient is almost close to zero. This is robust evidence indicating that 7 or 12 are the optimal cluster numbers to depict the local diffusion pattern of thalamic nuclei. The group thalamus atlas validation indexes can be seen in supplementary sheet 2.

**Fig. 3.**
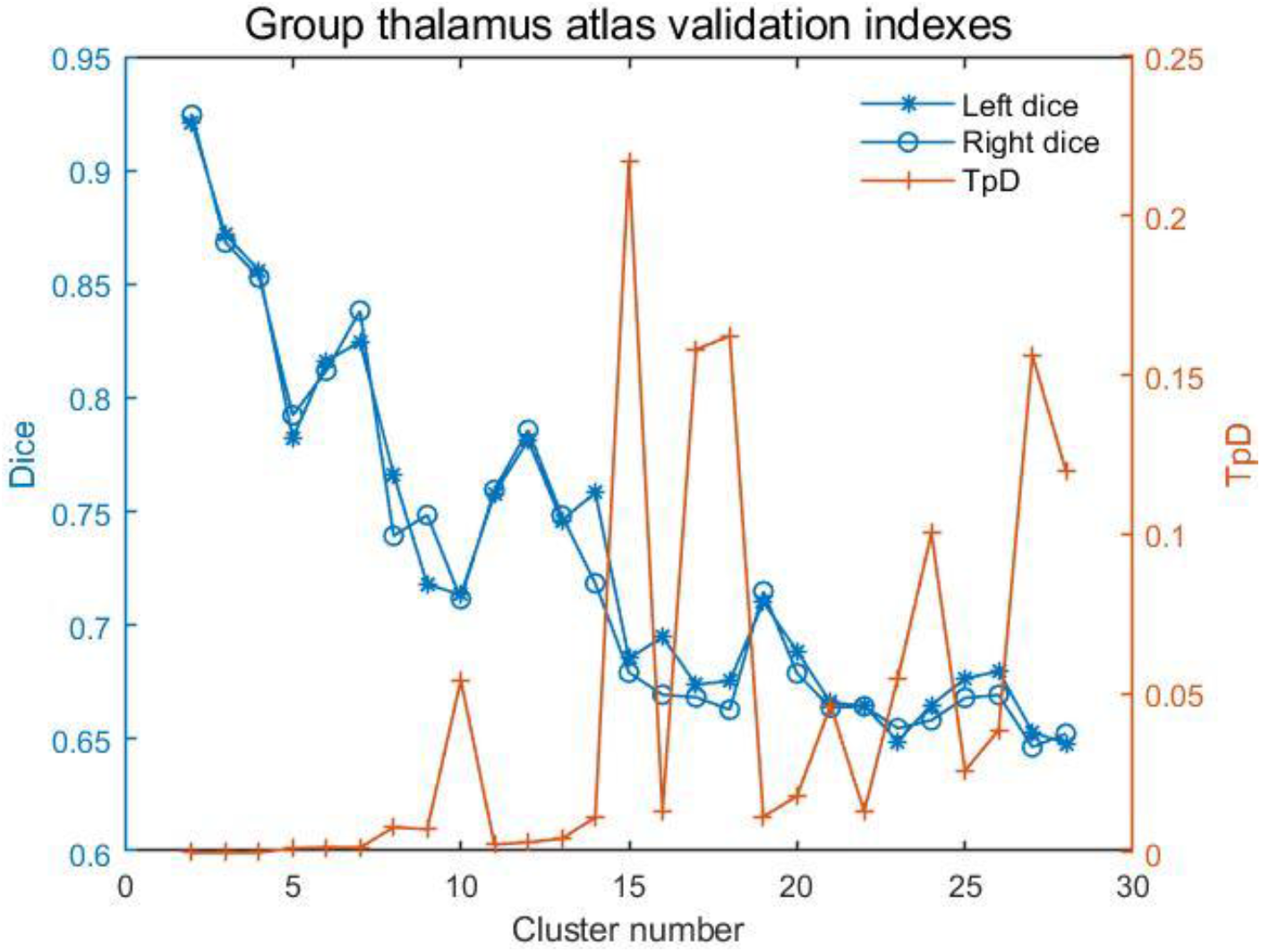
Validation indexes of group thalamus atlas. When cluster numbers are 7 and 12, the *Dice* coefficient shows acceptable local maxima in both left and right hemispheres. And the *Tpd* coefficient is almost zero. This indicates that 7 or 12 clusters are the intrinsic parcellation pattern of the thalamus.

ITK-SNAP ^83^ was used to visualize the parcellation pattern of the thalamus. In S.3, the thalamus is parcellated into 7 thalamic nuclei, and the parcellation pattern is consistent with the reliable ODF-based thalamus atlas construction study ^25^. Furthermore, higher data quality and a larger population allow higher-order ODF estimation to precisely and unbiasedly construct the thalamus atlas in tinier granularity. Thanks to the well-scanned and carefully defined HCP Unrelated 100 data sets, shown in Fig. 4, the 12-cluster thalamus atlas shows a significantly homologous parcellation pattern between two hemispheres and great consistency with thalamus anatomic structure. Besides, in this fine-grained group thalamus atlas, some subtle thalamic nuclei can be detected, such as dorsal anterior mediodorsal thalamic nucleus (MDda), dorsal posterior mediodorsal thalamic nucleus (MDdp), and ventral mediodorsal thalamic nucleus (MDv). As shown in Fig. 5(a), along with the growth of the probability threshold, the voxel number of all thalamic nuclei is evenly decreasing. In Fig. 5(b), the boundary voxels of thalamic nuclei gradually fade away with the increasing probability threshold, but the central area has been retained. This indicates that the nuclei boundaries of the group atlas reflect individual variation, while the central region represents population commonality.

**Fig. 4.**
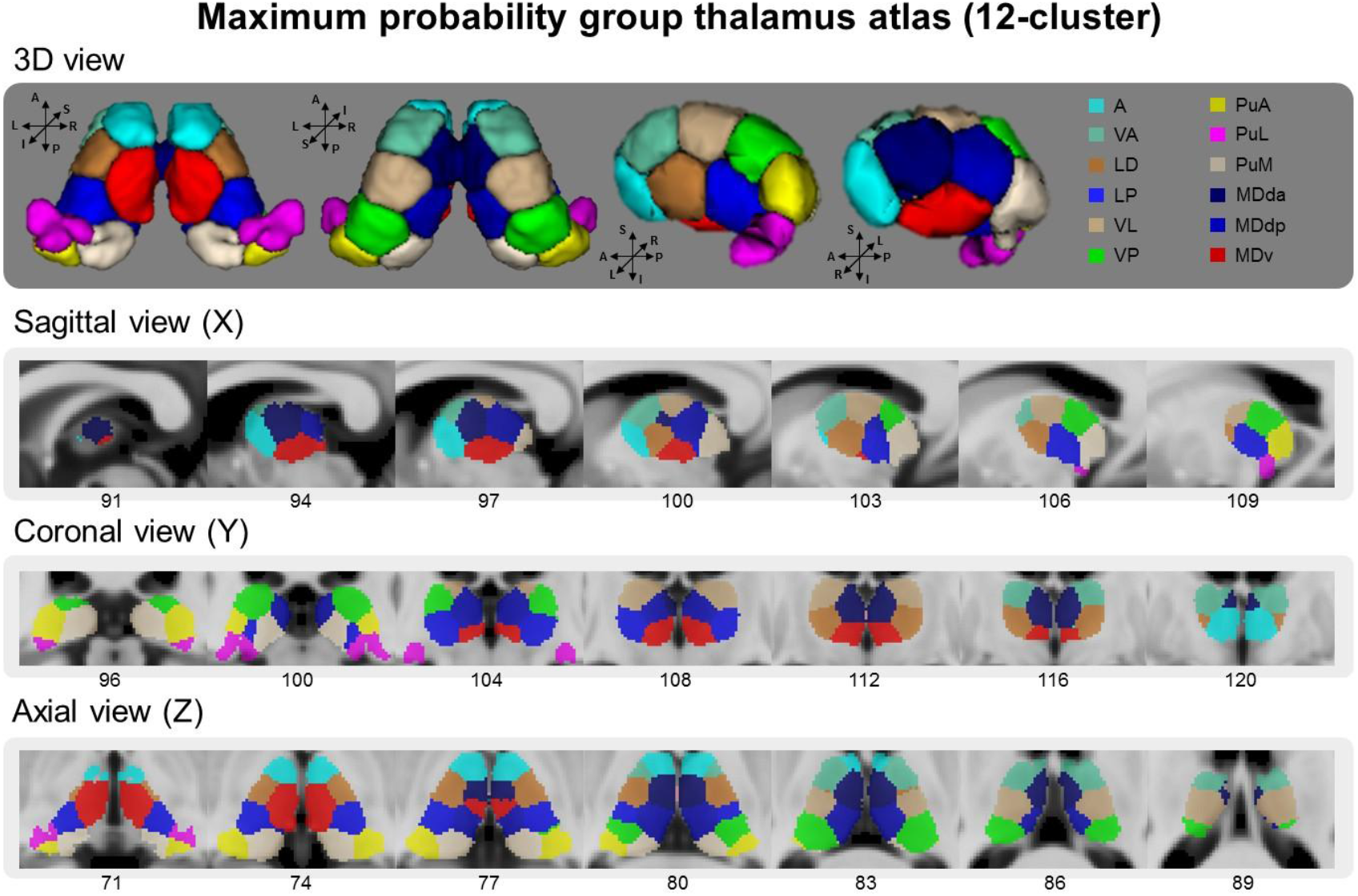
Maximum probability map of thalamus when cluster number is 12. In this group, thalamus atlas, the thalamus is parcellated into the anterior thalamic nucleus (A), ventral anterior thalamic nucleus (VA), lateral dorsal thalamic nucleus (LD), lateral posterior thalamic nucleus (LP), ventral lateral thalamic nucleus (VL), ventral posterior thalamic nucleus (VP), anterior thalamic pulvinar nucleus (PuA), lateral thalamic pulvinar nucleus (PuL), medial thalamic pulvinar nucleus (PuM), dorsal anterior mediodorsal thalamic nucleus (MDda), dorsal posterior mediodorsal thalamic nucleus (MDdp), ventral mediodorsal thalamic nucleus (MDv). The thalamus atlas shows strong homology between two hemispheres in a tinier granularity. Compared to 7-cluster, the 12-cluster thalamus atlas is more consistent with the actual thalamic anatomical structure.

**Fig. 5.**
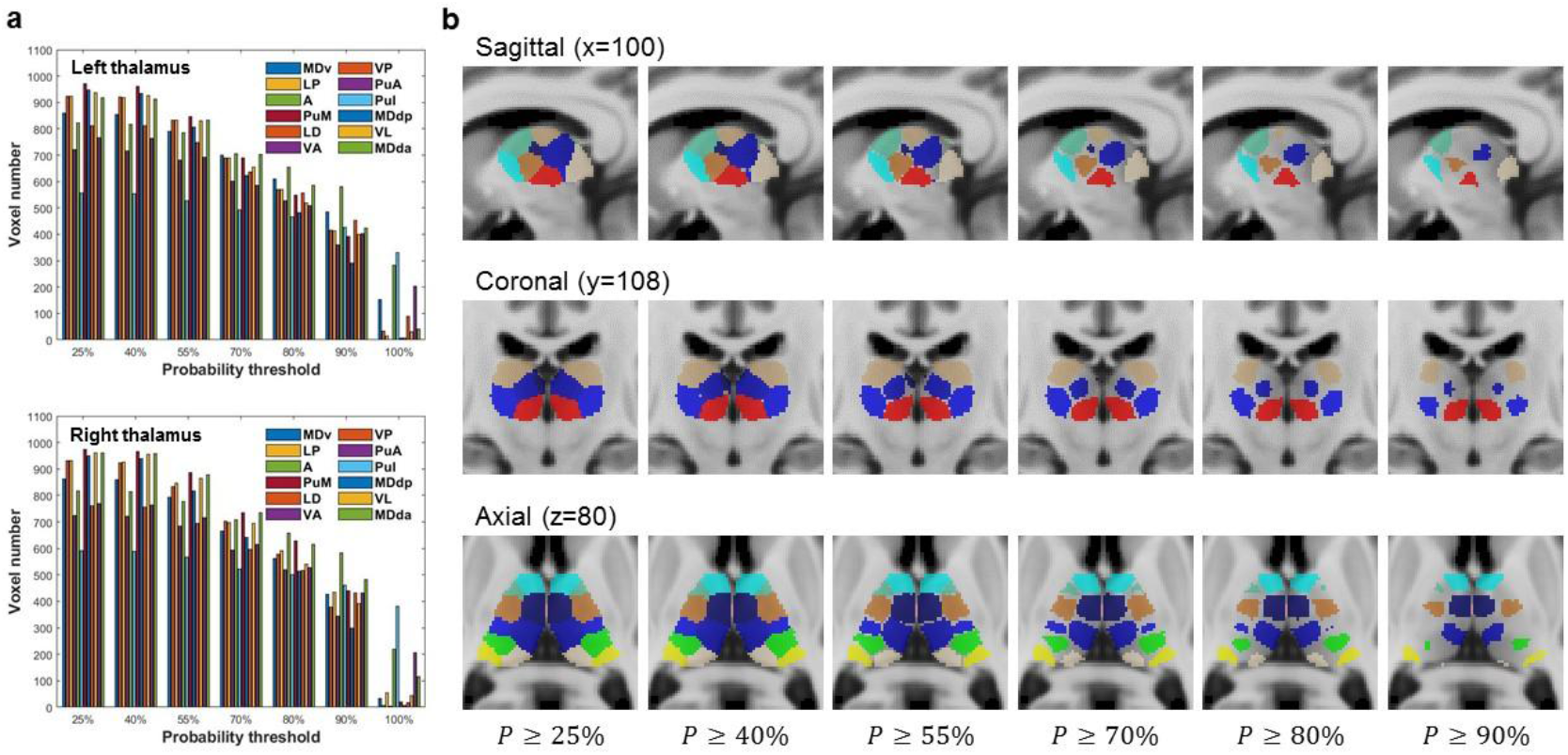
Probability map of thalamus when cluster number is 12. (a) Voxel number gradually reduces along with the increase of probability threshold of group atlas. (b) When the probability threshold rises, boundary voxels gradually fade, but the central area has been retained. This indicates that boundary voxels of the group thalamus atlas depict the individual variation, while the central region represents the population commonality.

### Individualized thalamus atlas

Based on the two potential optimal cluster numbers, the individualized thalamus atlas was constructed by three different individualization strategies, including group prior guided (PG), group atlas registration (GR), and single-subject clustering (CL). And the feasibility and robustness of these three pipelines were evaluated by four HCP data sets. The group probability threshold and 10-fold validation accuracy of each subject can be seen in Fig. 6(a) and S.5(a). It could be proved that each individualization model is guided by the high-confidence population commonality. Besides, the individualization models are highly generalizable and powerful to accurately classify the voxels into corresponding clusters. The silhouette coefficient was used to quantify the clustering goodness, that is, cohesion within and separation between clusters. As seen in Fig. 6(b) and S.5(b), in four HCP data sets, these three individualization strategies based on individualized thalamus atlases provide almost identical clustering goodness, and our PG (red) one has higher clustering goodness than GR (blue) and CL (yellow) (p<0.001). The clustering goodness is so close because the cluster labels of central voxels in thalamic nuclei are identical between the three individualization pipelines. In contrast, a small proportion of boundary voxels provides the main variation. But it can also be proved that our PG individualized thalamus atlas has the best clustering compared to GR and CL.

**Fig. 6.**
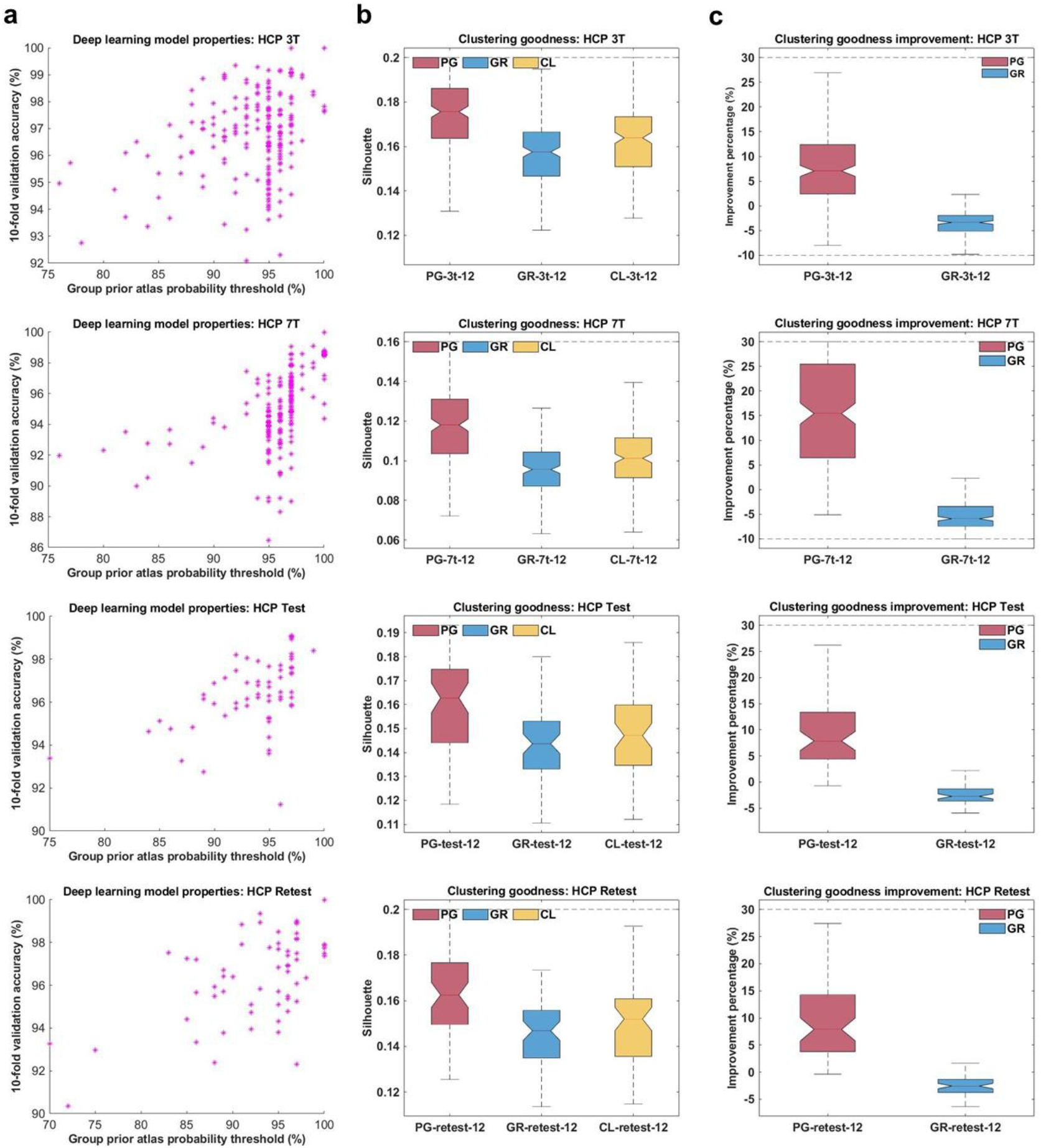
Validation indexes of individualized thalamus atlas when cluster number is 12. (a) Probability threshold of group prior atlas and 10-fold validation accuracy of prior-guided individualization model of all subjects. In each data set, both of 10-fold validation accuracy and probability threshold of the group prior atlas are significantly high. (b) Clustering goodness of three different individualization strategies driven thalamus atlases. In each data set, the prior-guided individualized thalamus atlas has the highest clustering goodness. (c) Improvement of clustering goodness of prior-guided and group atlas registration individualized thalamus atlas compared to single-subject clustering. In each data set, the improvement of prior-guided individualized thalamus atlas is positive, while group atlas registration is negative. PG: group prior guided. GR: group atlas registration. CL: single-subject clustering.

Small changes often make a huge difference. We quantitively examined the clustering goodness improvement of PG and GR individualized thalamus atlas compared to CL. As seen in Fig. 6(c) and S.5(c), in four HCP data sets, the PG (red) individualized thalamus atlas has significant growth in clustering goodness compared to CL. In contrast, the GR (blue) does have weaker performance, as reported in previous studies ^62^. Besides, the improvement of the 12-cluster individualized thalamus atlas is conspicuously more significant than the 7-cluster (seen in Fig.6(c) and S.5(c)). This is undoubted evidence to demonstrate that our PG individualized thalamus atlas has conspicuous better performance to depict the individual specificity at a tinier granularity level. When looking into the 3T-7T and test-retest validation, the 7T data gives a more considerable improvement than 3T, and the test-retest improvement is at a similar level. This is because more cluster numbers and higher spatial resolution provide more variation of boundary voxels to capture the individual specificity. The individualized thalamus atlas validation indexes can be seen in supplementary sheet 3 and 4.

We randomly chose three subjects to visually compare these three individualized thalamus atlases at 12-cluster parcellation granularity. As seen in Fig. 7, in different magnetic field strengths and scanning batches, the PG (first row) individualized thalamus atlas can more keenly and robustly detect the boundary of thalamic nuclei compared to GR (second row) and CL (third row).

**Fig. 7.**
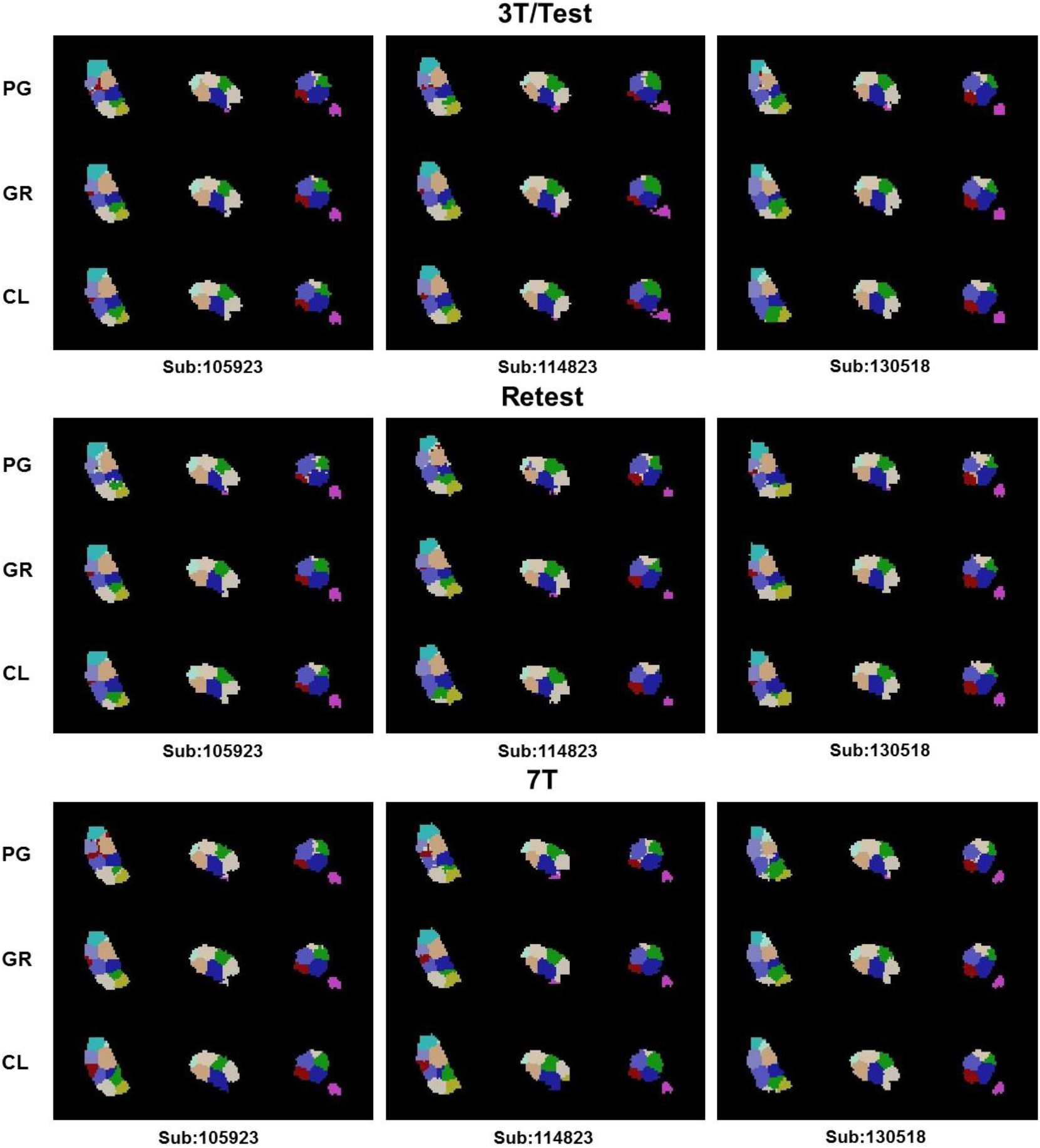
12-cluster individualized thalamus atlas constructed by three different individualization strategies. We randomly chose three subjects (105923, 114823, 130518) simultaneously existing in four HCP data sets and constructed their individualized thalamus atlases with three individualization strategies. In different magnetic field strengths and scanning batches, the prior-guided thalamus atlas can better detect the nuclei boundaries than the other two individualized thalamus atlases.

Based on the MRI data offered by Allen brain dataset, we constructed the individualized thalamus atlases by PG, GR, and CL individualization strategies. We compared them with the golden standard thalamic nuclei boundary, which was manually labeled according to Nissl staining. As shown in Fig. 8, compared to GR and CL, the PG individualized thalamus atlas is most consistent with the manually labeled histological atlas, which is regarded as the golden standard of thalamic parcellation. Besides, the PG individualized thalamus atlas can capture some subtle thalamic structures while GR and CL cannot.

**Fig. 8.**
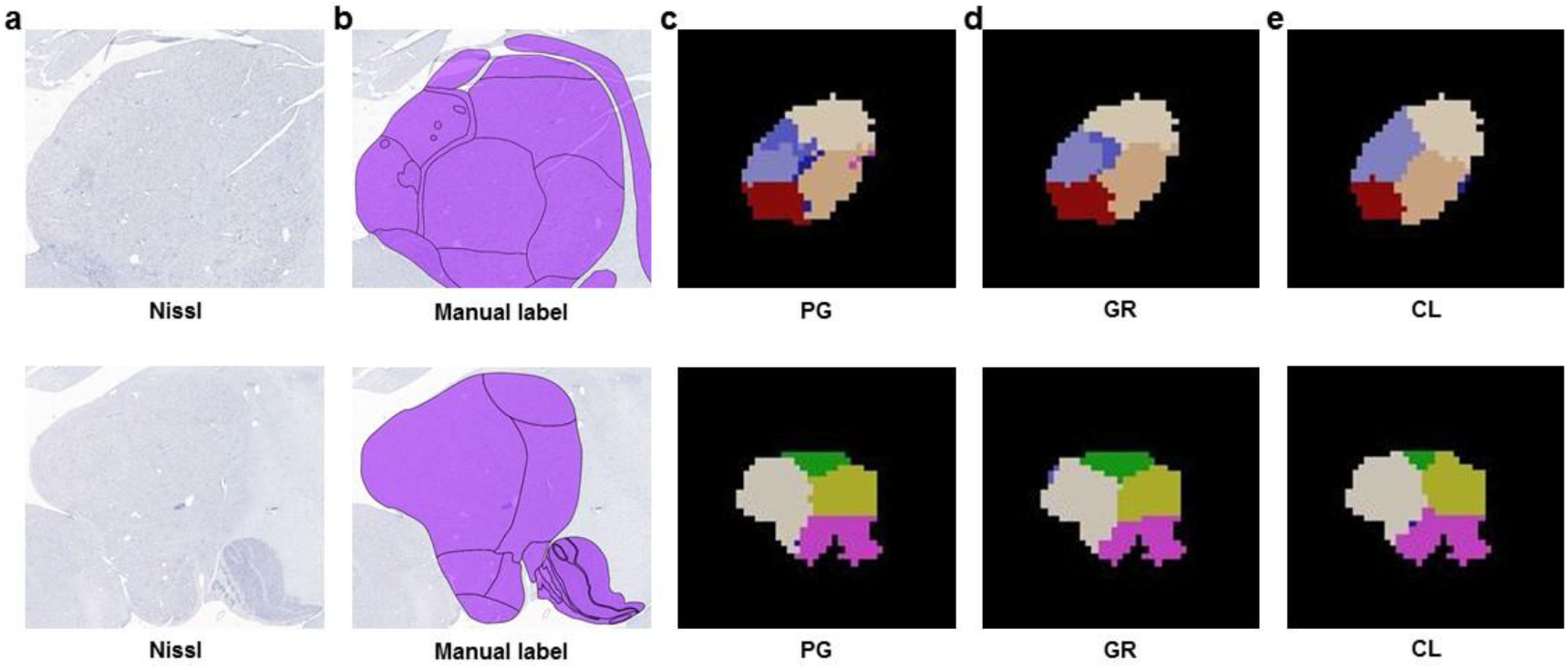
Histological validation of 12-cluster individualized thalamus atlas. (a) Nissl staining in the thalamic region. (b) Manually labeled thalamic nuclei boundaries. (c) Prior-guided individualized thalamus atlas. (d) Group atlas registration individualized thalamus atlas. (e) Single-subject clustering individualized thalamus atlas. Compared to GR and CL, the PG individualized thalamus atlas can more accurately detect the thalamic nuclei boundaries. Besides, the PG individualized thalamus atlas can precisely capture the subtle thalamic structure while both GR and CL cannot.

## Discussion

Using the high-quality HCP Unrelated 100 data set, we constructed the group probability thalamus atlas based on local diffusion features of thalamic nuclei. Then, we extracted the high-confidence group atlas as the prior guidance to construct the individualized thalamus atlas for each new subject by deep learning-based semi-supervised classification model. We found that the prior-guided individualized thalamus atlas is more capable of depicting the individual specificity and detecting the thalamic nuclei boundaries than single-subject-based clustering and group atlas registration. And in finer parcellation granularity, the prior-guided individualized thalamus atlas provides a more conspicuous performance than the coarse-grained parcellation pattern. Besides, through the 3.0T-7.0T and test-retest validation, our prior-guided individualized thalamus atlas construction pipeline is robust and repeatable to different magnetic field strengths and scanning batches. In addition, examined by the Allen brain data set, our prior-guided individualized thalamus atlas has the highest histological consistency compared to single-subject clustering and group atlas registration.

Here, we proposed a prior-guided individualization paradigm to solve the issue of objectively combining the population commonality and individual specificity when constructing the brain atlas. By partly aligning the central area of the group atlas into the native space and generating the individualization model for each specific subject, the prior-guided individualization approach can effectively overcome the registration error across different spaces and provide an individualized atlas with significantly high personality. Besides, the prior guidance is generated by the canonical high-confidence group atlas and retained in the native space, offering the individualized brain atlas convincing population commonality. In contrast to group atlas registration, the single-subject-based approaches construct the individualized atlas using individual characteristics in the native space of the subject, which can provide higher individual specificity. However, both group atlas registration and single-subject-based individualized atlas construction pipelines roughly share the same parcellation parameters for each subject. This must have introduced group standard parameters constraints into individual atlases. To solve this, based on the deep learning algorithm, our prior-guided individualization pipeline constructs a personalized individualization model in the native space for each subject, allowing a chance to more accurately depict the individual specificity.

Compared to previous microstructural thalamic parcellation ^25,46,47^, the 12-cluster thalamus atlas provides a more fine-grained parcellation pattern of thalamus nuclei based on a higher-order ODF model. And the 12-cluster thalamus atlas is significantly consistent with the thalamic anatomic structure and homologous between hemispheres. Besides, some subtle thalamic structures could be detected by our 12-cluster thalamus atlas, which may reveal the more detailed intrinsic microstructural organization pattern of the thalamus. Guided by the high-probability group prior atlas, we developed an automatic individualized thalamus atlas construction pipeline based on a deep learning algorithm. Existing individualized thalamus atlases are constructed by either single-subject-based clustering ^25,47^, white matter tractography quantification ^24^, or multi-subject-based group atlas registration ^66^. Compared to these approaches, our prior-guided individualization pipeline is the first attempt to construct the microstructural individualized thalamus atlas in the native space of the subject by combining group prior atlas and individual diffusion properties. The HCP data set has shown that an individualized thalamus atlas combing population commonality and individual specificity is superior to group atlas registration and single-subject clustering in depicting the individual thalamic nuclei boundaries.

In the current study, the well-scanned and carefully defined HCP Unrelated 100 data set was used to construct the group thalamus atlas. However, a larger population could be adopted to construct a more reliable group thalamus atlas, which provides a more persuasive group prior guidance. Thanks to the multiple *b*-values diffusion-weighted scanning dMRI data, the 8th-order ODF model can be reconstructed to exquisitely depict the detailed local diffusion properties of thalamic nuclei by reliable estimation algorithm ^44,75,76^. But in practice, longer scanning time and higher economic cost may limit the collection of multiple *b*-values dMRI data. To solve this, the single *b*-value based diffusion tensor model ^84^ or constrained spherical deconvolution ^43^ models with lower complexity could be adopted to quantify the diffusion properties of thalamic microstructure. Besides, the spatial resolution of HCP 3T and 7T dMRI is 1.25mm isotropic and 1.05mm isotropic, respectively. The relatively high spatial resolution provides sufficient voxels to train an accurate model and depict the individual specificity. However, when the spatial resolution becomes lower, such as the clinical 3T dMRI with 2.0mm isotropic spatial resolution, the efficiency of the deep learning model may decrease due to the insufficient train set. Also, the boundary voxels of thalamic nuclei may be too few to depict the individual variation. Here, some concise multiple classification models, such as support vector machine ^85^ and random forest ^86^, or transfer learning approach ^87,88^ could be applied to reduce the dependence on data volumes and the feature dimensions. In addition, super-resolution techniques ^89,90^ are clever alternatives to increase the train set’s information capacity by enhancing the spatial resolution feature maps.

In conclusion, we proposed a new prior-guided individualization paradigm for thalamic parcellation, providing a robust and more powerful ability to depict the individual specificity and detect the nuclei boundaries. Our prior-guided individualized thalamus atlas construction strategy can offer a reliable paradigm for individualized brain atlas construction. Further, we plan to construct the individualized brain atlas using clinical MRI data and pursue clinical validation. Besides, based on this individualized thalamus atlas construction pipeline, it is worth assessing the performance of prior-guided individualized brain atlas, such as cognition, behavior, aging, development, and disease. Also, the prior-guided clinical individualized brain atlas may provide excellent accuracy in disease diagnosis, neurosurgery localization, and prognosis prediction.

## Acknowledgments

This work was partially supported by Science and Technology Innovation 2030 - Brain Science and Brain-Inspired Intelligence Project of China (Grant No. 2021ZD0200200), the Natural Science Foundation of China (Grant Nos. 82072099, 91432302, and 31620103905), the Strategic Priority Research Program of Chinese Academy of Sciences (XDB32030200), National Key R&D Program of China (Grant No. 2017YFA0105203), the Youth Innovation Promotion Association, and the Beijing Advanced Discipline Fund.

## Supplementary Material

It is shown in supplementary figures and supplementary sheets.

## Data and Code Availability

The HCP S1200 data is available at https://humanconnectome.org. The FSL can be downloaded at https://fsl.fmrib.ox.ac.uk/fsl/. The SPM can be downloaded at https://www.fil.ion.ucl.ac.uk/spm/.

The MRtrix3 can be downloaded at https://www.mrtrix.org. The ITK-SNAP can be downloaded at http://www.itksnap.org. The source code of this work will be made public after the paper is published.

