## Supplementary material for "Prior-guided Individualized Thalamic Parcellation based on Local Diffusion Characteristics using Deep Learning": Supplementary figures.docx

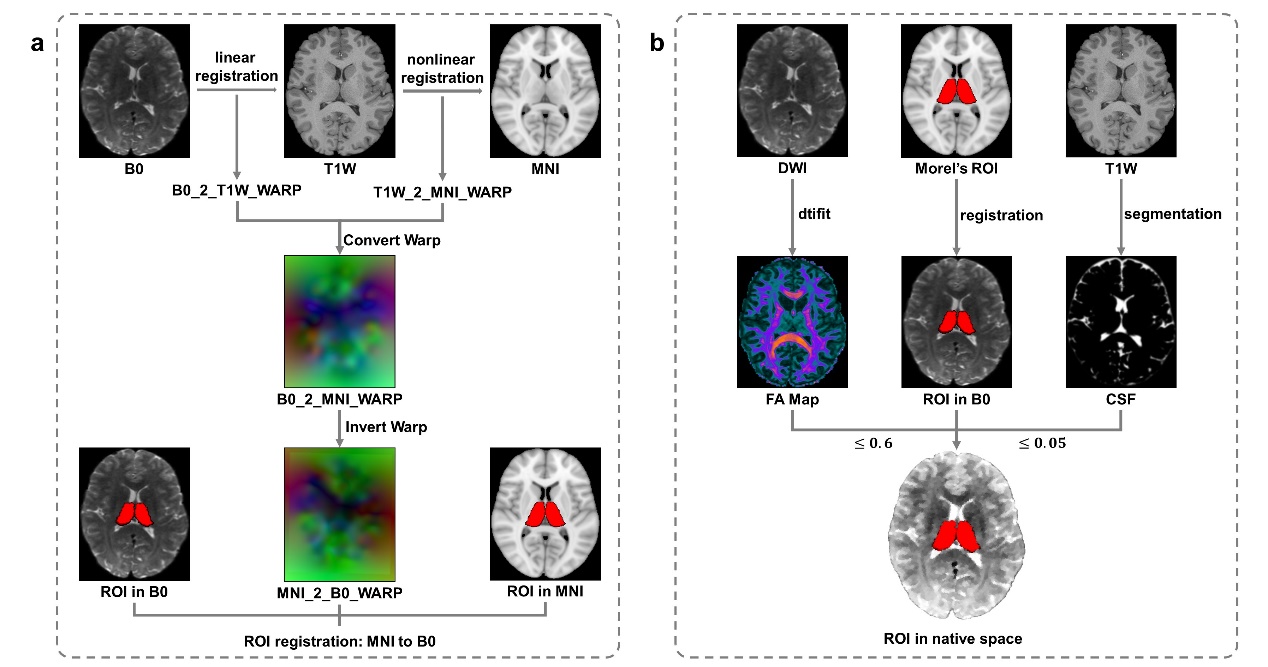

**S.1 | ROI registration and postprocessing.**

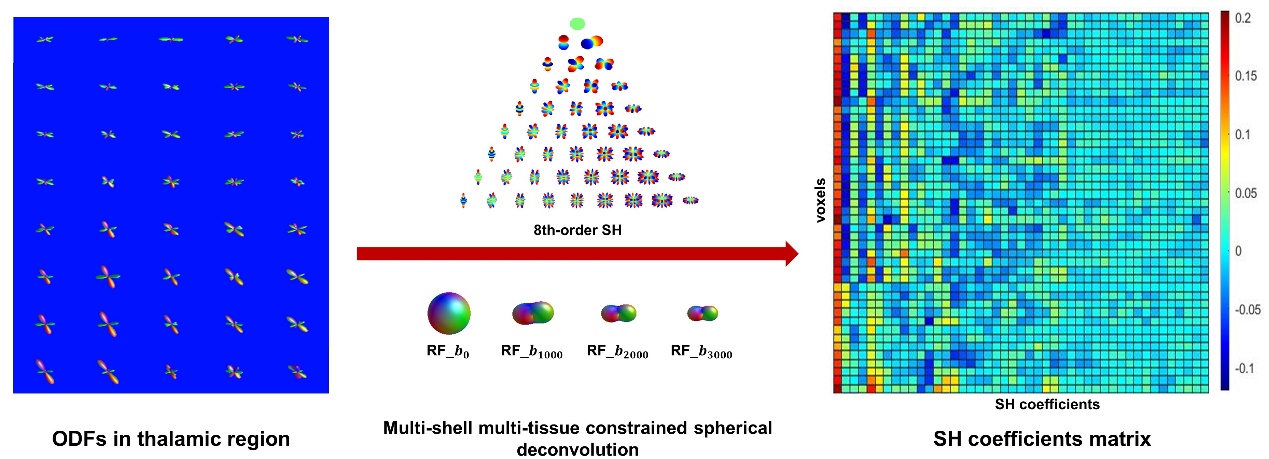

**S.2 | ODF estimation.**

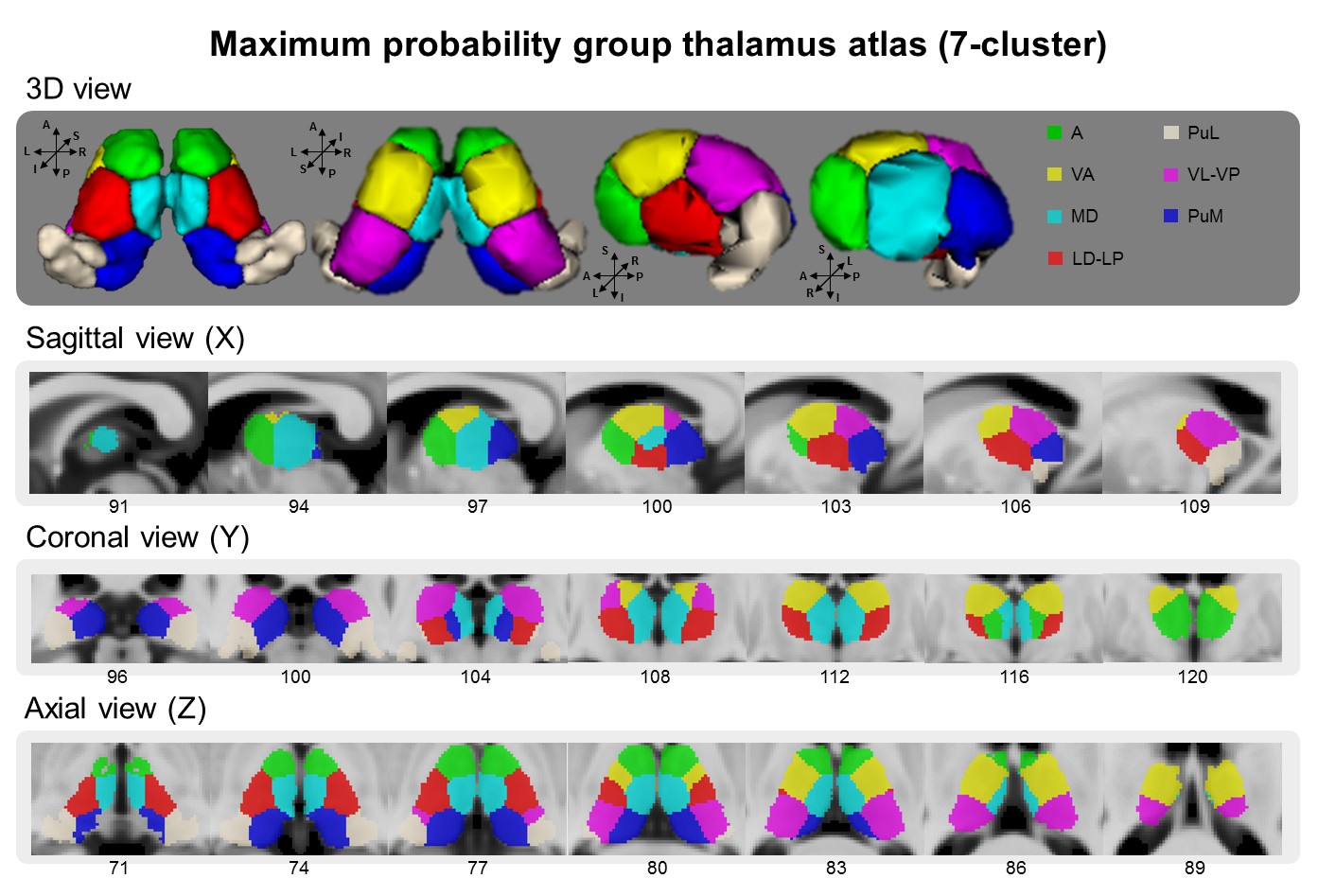

**S.3 | Maximum probability map of thalamus when cluster number is 7.**

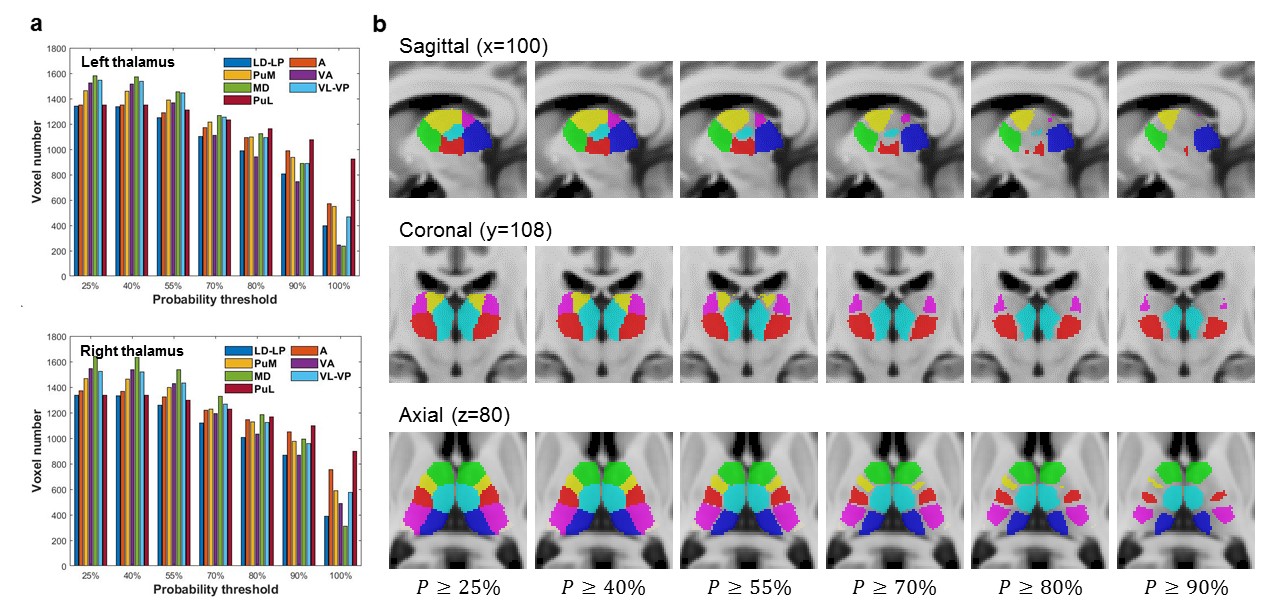

**S.4 | Probability map of thalamus when cluster number is 7.**

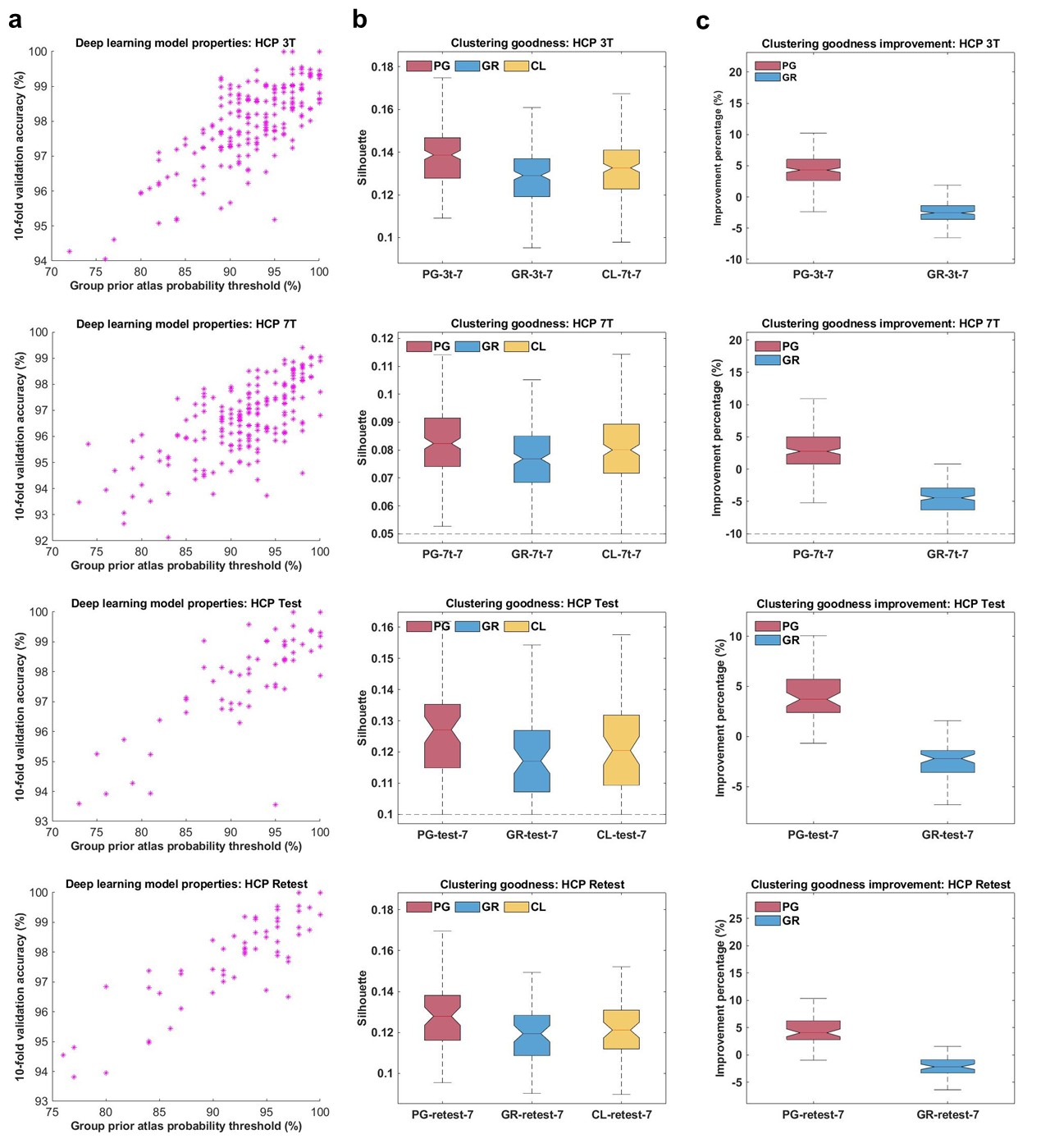

**S.5 | Validation indexes of individualized thalamus atlas when cluster number is 7.**
